# Trapalyzer: A computer program for quantitative analyses in fluorescent live-imaging studies of Neutrophil Extracellular Trap formation

**DOI:** 10.1101/2022.12.14.520407

**Authors:** Michał Aleksander Ciach, Grzegorz Bokota, Aneta Manda-Handzlik, Weronika Kuźmicka, Urszula Demkow, Anna Gambin

## Abstract

Neutrophil extracellular traps (NETs), pathogen-ensnaring structures formed by neutrophils by expelling their DNA into the environment, are believed to play an important role in immunity and autoimmune diseases. In recent years, a growing attention has been put into developing software tools to quantify NETs in fluorescent microscopy images. However, current solutions require extensive training data sets, are difficult to use for users without background in computer science, or have limited capabilities.

In this work we present Trapalyzer, a computer program for an automatic quantification of NETs in terms of their area and an approximation of their number. In addition, Trapalyzer counts neutrophils at different stages of NET formation, and is the first tool that makes this possible without extensive training data sets. We validate our approach on a publicly available benchmark data set and apply it in a neutrophil-bacteria co-culture experiment. The software and usage tutorials are available at https://github.com/Czaki/Trapalyzer.

## 1 Introduction

Neutrophils are the most abundant group of white blood cells in humans. They are often described as the organism’s “frontline soldiers”, responsible for fighting pathogens during the initial stage of infection [16, 12]. One of their fighting strategies is the formation of Neutrophil Extracellular Traps (NETs), web-like structures formed from the cells’ DNA which ensnare and putatively kill microbes [6, 1]. NETs help to fight infections, but may also harm the host by damaging surrounding tissues and promoting inflammation. Research shows that excessive or insufficient formation of NETs plays a role in a number of diseases, including periodontitis, thrombosis, and arthritis [7, 3]. A better understanding of the dynamics of NET formation may lead to improved diagnostics and treatment of those diseases.

There are numerous experimental methods of NET quantification [10, 13, 3]. A popular technique is to stain the extracellular DNA with a cell-impermeable fluorescent dye, e.g. SYTOX™ Green, and count the number of NETs by eye under a fluorescent miscroscope. With this staining, NETs appear as large, fibrous or cloud-like, often irregular structures [3]. Adding a cell-permeable dye, such as Hoechst 33342, makes it possible to visualize both NETs and cells that did not release the traps and estimate the rate of NET formation.

In recent years, there has been a growing interest in developing computational methods of NET quantification to make it more replicable and objective, while at the same time less laborsome and time-consuming [19]. A number of programs has been released, either based on machine learning algorithms, including convolutional neural networks and support vector machines [9, 11, 14], or digital image processing techniques, including image thresholding and classification of regions of interest (ROIs) based on features such as area or circularity [17, 15, 5]. Modern machine learning-based methods are capable of quantifying not only NETs, but also cells at certain stages of NET formation, giving a greater insight into the dynamics of this process [14].

Currently, however, both groups of tools have their drawbacks which limit their usability. Computer programs based on machine learning require laborious manual preparations of large training data sets and are often difficult to use for users without a computer science background. On the other hand, the currently available tools that are free from those requirements quantify NETs, but not the numbers of neutrophils at different stages of NET formation. One of the reasons for this situation is that they are arguably more difficult to develop. While machine learning algorithms, given a manually annotated data set, are able to figure out the crucial steps of image annotation by themselves, tools based on digital image processing techniques need an explicit, human-designed algorithm for this task. Developing such an algorithm requires an in-depth expert knowledge of the analyzed process and dedicated studies on how to mathematically describe and distinguish different cell morphologies.

### Our contribution

In this work, we present Trapalyzer, a plug-in for the PartSeg software [4] for the analysis and annotation of fluorescent microscopy images of neutrophils double-stained with a cell-permeable and a cell-impermeable fluorescent DNA dye. Our software extends the capabilities of the currently available tools by quantifying more stages of NET formation without the need for large training data sets. This has been made possible by extensive studies of fluorescent microscopy images by an interdisciplinary team composed of clinical scientists, statisticians, and computer scientists, which have resulted in a small set of ROI features that characterize the stages, and a scoring system that uses those features to classify cells. Trapalyzer also detects autofluorescence artifacts in the green channel, making NET quantification more reliable and robust.

Trapalyzer offers two modes of analysis: an interactive session and a batch processing mode. The interactive session allows the user to set the program’s parameters and visualize the annotation, while the batch processing mode can be used to process multiple images in a single run and save the results in a convenient Excel spreadsheet. The user can specify the information to be computed, both image-wise (such as the number of neutrophils at a given stage of NET formation, the percent of image area covered by NETs, or the quality of annotation) and ROI-wise (such as the size of each ROI, its bounding box, or assigned class).

We validate our approach on a publicly available benchmark data set published in [14] and show that it attains a similar performance to convolutional neural networks using just a fraction of the training data set. We then show how Trapalyzer can be applied to an experiment on the dynamics of neutrophil-*E. coli* bacteria interactions, where we study the cells’ progression through the stages of NET formation. The results of this experiment agree with observations made for individual cells by other authors [18].

Trapalyzer is designed with an emphasis on software ergonomy and ease of use. The plug-in requires no installation other than downloading and placing in the PartSeg’s directory and is accompanied with easy to follow tutorials available on the project’s website. The tutorials guide the users through a stepby-step procedure to tune the program’s parameters and configure its output. This allows the users to easily learn how to use the software and apply it to their own experiments even if they have no background in computer science, giving Trapalyzer the potential to be routinely used in laboratories researching diverse aspects of NET formation.

## 2 Background

The process of NET formation has been recently studied on a cellular level by the means of high-resolution time-lapse microscopy [18]. The authors have observed that this process progresses through a sequence of stages, shown schematically in Fig. 1. Intact, unstimulated neutrophils are characterized by a lobulated nucleus with spatially heterogeneous condensation of chromatin. The onset of NET formation is marked by the disassembly of the actin cytoskeleton and the formation of plasma membrane microvesicles containing cytosolic components. Next, the microtubule and vimentin cytoskeletons undergo disassembly and remodeling, after which the neutrophils’ chromatin undergoes decondensation, with its fluorescent staining becoming spatially homogeneous. During and after chromatin decondensation, the nucleus loses its lobulation and becomes partially or fully rounded, after which the nuclear envelope ruptures, causing a leakage of the DNA into the cytoplasm. At a similar time, the plasma membrane gradually increases its permeability, causing membrane-impermeable markers to enter the cell. Finally, the plasma membrane ruptures, releasing the genetic material to the environment.

**Figure 1:**
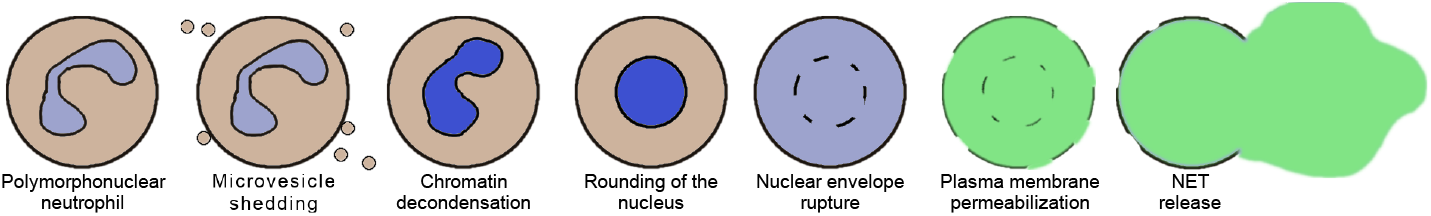
A schematic representation of the stages of Neutrophil Extracellular Trap (NET) formation, based on [18]

The transitions between different stages take different amounts of time. The decondensation of chromatin and rounding of the nucleus are gradual processes. After their completion, the neutrophil typically spends some time in the rounded nucleus state before a rapid disruption of the nuclear envelope and release of chromatin into the cytoplasm. The permeabilisation of the plasma membrane is also gradual and typically completes after the nuclear envelope rupture, after which the neutrophil spends some time with its DNA fully stained by cell-impermeable DNA markers. After that, NET release occurs in some, but usually not all neutrophils, and may be either rapid or gradual [18].

The complex nature of the process of NET formation poses a substantial difficulty in developing software tools to analyze it. Furthermore, which stages are suitable for automatic quantification depends on the experimental method used. For example, microvesicle shedding is visible using high-resolution differential interference contrast microscopy, but not in fluorescent microscopic images of stained DNA. It is a challenging task to pinpoint distinct cell morphologies that can be rigorously quantified, provide their mathematical characterization, and use it to develop an algorithm for an automatic image annotation.

The quantification of NETs poses some still unsolved challenges as well. The state of the art tools usually detect NETs as large ROIs with intense brightness in the extracellular DNA channel, sometimes taking low circularity as an additional criterion under an assumption that NETs have irregular boundaries [17, 15]. However, when the DNA is stained with a green dye, such as SYTOX™ Green, autofluorescence on the green channel may cause artifacts in the images [2]. Such artifacts can have similar areas and shapes to NETs, posing yet another challenge for an automated image annotation tool.

## 3 Methods

To establish the quantifiable classes of ROIs for NET formation studies, we have performed a neutrophil killing assay of *Escherichia coli* bacteria. To verify our conclusions and to assess the performance of our approach we have downloaded a benchmark set of images in which neutrophils were incubated without bacteria and NET formation was induced by various chemical stimuli.

### Reagents

Roswell Park Memorial Institute (RPMI) 1640 medium, HEPES, SYTOX™ Green, and Hoechst 33342 were purchased from Thermo Fisher Scientific (Waltham, USA). LB broth was purchased from Sigma Aldrich (St Louis, MO, USA).

### Preparation of blood neutrophils

Neutrophils were obtained from peripheral blood of one healthy blood donor. Blood sample was purchased at Local Blood Donation Centre and according to local regulations, the blood donor enabled blood donation center to sell their blood samples for scientific purposes and the consent of bioethical committee was not required. Blood was collected into a citrate tube and processed within 2 hours from collection. Neutrophils were isolated using density gradient centrifugation followed by polyvinyl alcohol sedimentation, exactly as described in [8]. Isolated neutrophils were suspended in RPMI 1640 medium with 10 mM HEPES (RH).

### Preparation of bacteria

*Escherichia coli* (American Type Culture Collection(ATCC) 25922 strain) were grown overnight in LB broth with shaking.

In the morning, an aliquot of bacterial culture was taken, diluted 100 x in a fresh LB medium and grown for subsequent 2-3 hours. Subsequently, bacterial cultures were washed and resuspended in RH medium.

### Co-culture of neutrophils with bacteria

Neutrophils were seeded into the wells of 48-well plates at the density of 2 × 10^4^ cells/well and allowed to settle for 30 minutes at 37°C, 5% CO2. Subsequently, *E. coli* was added into the appropriate wells at the multiplicity of infection of 4 or 1 (*E*.*coli* : neutrophil).

Neutrophils incubated without bacteria were used as a control group. A technical duplicate for each condition was prepared. For each intended timepoint (t=0, 60, 90, 120, 180 minutes), a separate 48 well plate was prepared. The plates were centrifuged for 5 minutes at 250 g to allow the contact of bacteria with neutrophils. The plates were incubated at 37°C, 5% CO2 for a specified time and then the samples were stained with SYTOX™ Green (100 nM) and Hoechst 33342 (1.25 *μ*M) for 10 minutes. Four images of each well were taken with Leica DMi8 fluorescent microscope equipped with a 10× magnification objective (Leica, Wetzlar, Germany). Overall, 120 images have been obtained.

### Benchmark data set

A benchmark data set of images of neutrophils and NETs stained with SYTOX™ Green and Hoechst 33342, published in [14], was downloaded from https://github.com/krzysztoffiok/CNN-based-image-analysis-for-detection-and-quantification-of-neutrophil-extracellular-traps on May 19, 2019. For evaluation of Trapalyzer’s accuracy, we have selected the validation set in file `large_validation_set.zip`, subdirectory `xml_pascal_voc_format/images/oryg`. The validation set consists of 57 images. Manual annotations of the images were accessed in subdirectory `xm l_pascal_voc_format/annotations/oryg`. Annotations in xml files were handled using the `lxml` library of the Python 3 programming language. For tuning of Trapalyzer’s parameters, additional 10 images were selected from the file `original_uncompressed_images_with_pascalvoc_annotations.zip`, rescaled to match the dimentions of the validation set images and converted to the TIFF format using the convert program from the `ImageMagick` suite.

## 4 Results

### 4.1 Quantifiable stages of NET formation

In order to pinpoint the stages of NET formation that are be suitable for quantification using an automated algorithm, we first analyzed manually a set microscopic images from a neutrophil-*E. coli* co-cultures.

#### Stages of NET formation identified with high-resolution time-lapse microscopy can be observed in low-resolution fluorescent microscopy

Most of the cells in the images taken at t=0 min. exhibited a typical appearance of unstimulated, polymorphonuclear neutrophils, without detectable signal in the extracellular channel (Fig. 2 A). In images taken between t=60 and t=120 min., we have observed cells which were visibly brighter and highly circular (Fig. 2 B). This morphology most likely corresponded to cells with a rounded nucleus. Individual cells exhibited this morphology in t=0 min. as well. We have also observed ROIs with larger areas, lower brightness, and cloud-like appearance, with no signal in the extracellular channel (Fig. 2 C). We assume that this morphology corresponded to cells with a ruptured nuclear envelope. We did not observe such cells in t=0 min. In images taken after t=60 min., and mostly in the later stages of the experiment, we have observed cells with cloudy appearance and detectable signal in the extracellular channel (Fig. 2 D). This morphology corresponds to cells with a permeabilized plasma membrane. The intensity of signal in the extracellular channel varied highly for those cells, indicating a gradual permeabilization, in agreement with [18].

**Figure 2:**
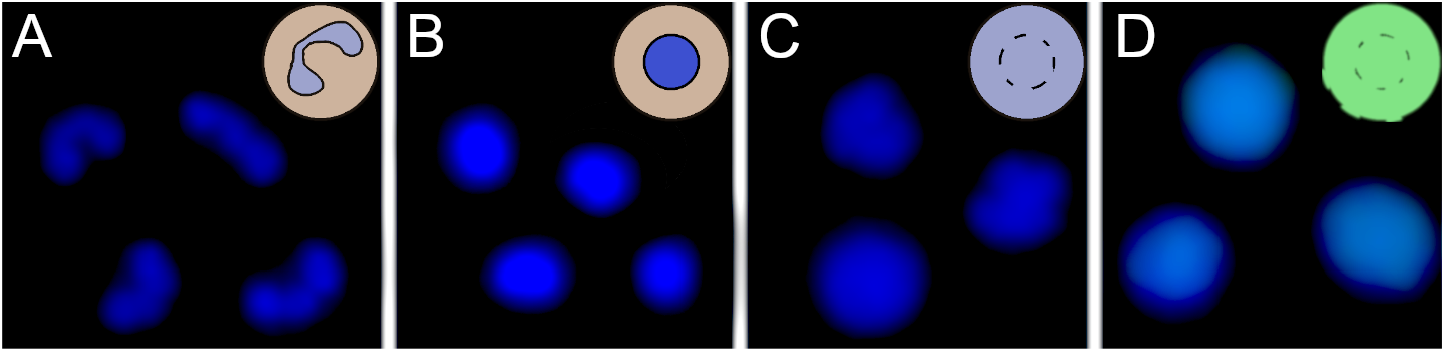
Neutrophils at different stages of NET formation visible in fluorescent microscopy with SYTOX™ Green/Hoechst 33342 double staining. The source images were taken as a part of the neutrophil-*E. coli* co-culture study. A) Polymorphonuclear (unstimulated) neutrophils; B) Neutrophils with rounded nucleus; C) Neutrophils with ruptured nuclear envelope; D) Neutrophils with permeabilized plasma membrane; E) A neutrophil extracellular trap.

#### >Not all stages of NET formation are suitable for an automatic quantification

Unstimulated neutrophils, cells with rounded nuclei, cells with ruptured nuclear envelopes and cells with permeabilized plasma membranes have distinct morphologies in fluorescent microscopy images, making them suitable for automatic detection by a software tool. On the other hand, neutrophils inbetween those stages, in particular neutrophils undergoing chromatin decondensation and nuclear rounding, are more difficult to classify. The stage of chromatin decondensation (between the onset of NET formation and nuclear rounding) does not seem to have a clear delineation from its surrounding stages, and does not appear to be suitable for automatic quantification, at least in fluorescent microscopy images of double-stained DNA.

#### Clumps of bacteria are an important class of ROIs in neutrophil killing assays

Starting from t=120 min., we observed clumps of bacteria, both in the total DNA channel and in the visible light (Fig. 3). In fluorescent light, they appeared as highly amorphous, low-brightness objects without well-defined edges. With the exponential growth of bacteria, those clumps become prevalent in t = 180 min., motivating the decision to include them as yet another class of ROIs.

**Figure 3:**
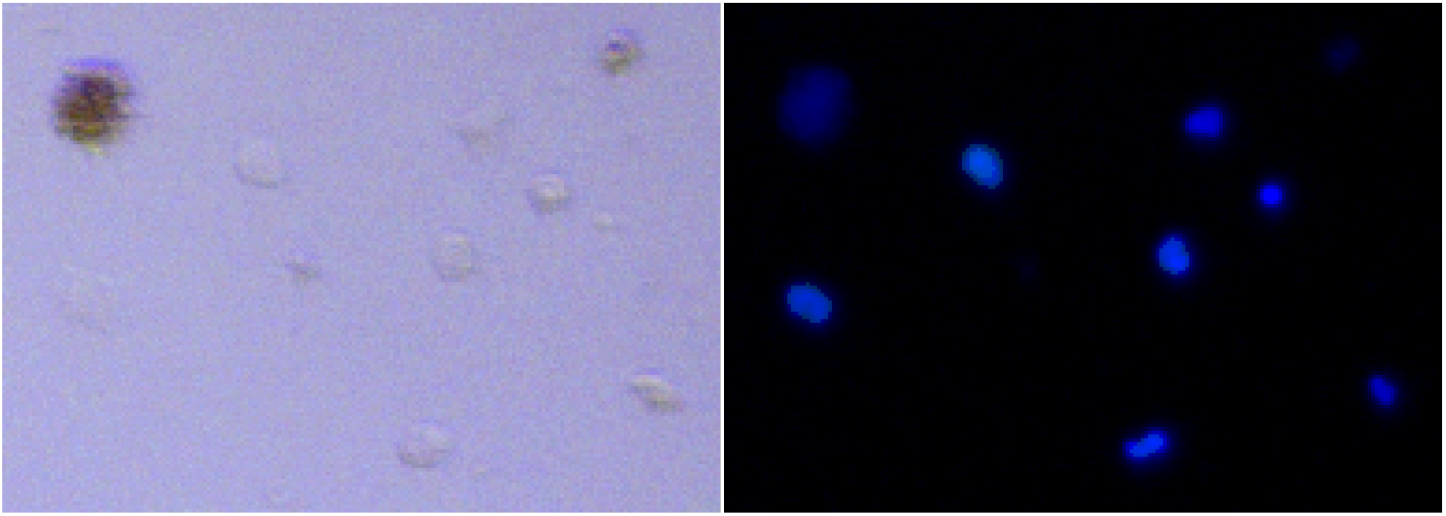
A characteristic appearance of a clump of bacterial cells (top left corner in both panels) in the visible light (left panel) and the fluorescent light (right panel). Both panels show the same fragment of an image.

#### Autofluorescene artifacts on the green channel are common

We have observed autofluorescence artifacts in approximately 30% of images. Most of them were small, and, apart from the possibility of being classified as NETs, did not pose other difficulties for automatic image annotations. However, some artifacts occupied extensive regions of the image, making the automated annotation unreliable (Fig. 4). This motivates the need for a score of annotation quality that reflects the area occupied by artifacts and other unclassified ROIs.

**Figure 4:**
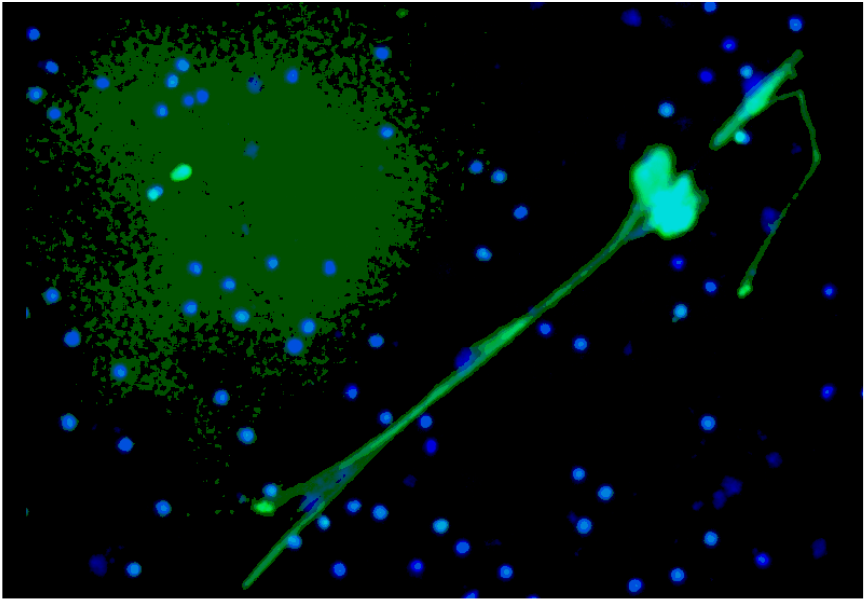
An autofluorescence artifact in the green channel after applying a median filter (top left corner), and a mature Neutrophil Extracellular Trap (diagonally throughout the image). Two emerging NETs and multiple neutrophils with and without permeabilized plasma membranes are located within the artifact.

### 4.2 ROI features at different stages

Based on the results presented in [18] and our analysis of fluorescent microscopy images, we consider seven classes of ROIs: unstimulated neutrophils; cells with decondensed chromatin and rounded nuclei; cells with ruptured nuclear envelopes; cells with permeabilized plasma membranes; neutrophil extracellular traps; clumps of bacteria; and autofluorescence artifacts. After fixing the set of ROIs that could potentially be quantified in fluorescent microscopy images of double-stained DNA, we looked for a minimal set of features that could be used to distinguish them.

#### Stages of NET formation have characteristic values of three ROI features

Polymorphonuclear neutrophils could be distinguished from other classes by a relatively small size and average brightness on the total DNA channel and the lack of signal in the extracellular DNA channel. Neutrophils with rounded nuclei could be distinguished from other ROIs by particularly high values of average brightness in the total DNA channel. Neutrophils with ruptured nuclear envelopes typically had larger areas than both previous classes. In some cases, they had similar areas to neutrophils with rounded nuclei, but could be distinguished from this class by lower average brightness. Neutrophils with permeabilized plasma membrane covered the same range of areas and brightness in the total DNA channel as the three previous classes. However, they could be easily distinguished from other cells as the only ones with detectable signal in the extracellular DNA channel.

#### Standard deviation of extracellular signal separates autofluorescence artifacts from NETs

Large areas and intense signal in the extracellular DNA channel distinguished NETs from cells, but not from autofluorescence artifacts. However, due to differences in chromatin density, the brightness of NETs was spatially non-homogeneous, while for autofluorescence artifacts it was mostly uniform, especially after applying a median filter. This was effectively captured by the standard deviation of brightness in the extracellular DNA channel. We did not observe an increase in classification accuracy when we included a measure of circularity used by other authors to detect NETs. On the contrary, NETs can be highly circular in shape (Fig. 2), especially when formed in the absence of bacteria [14].

#### Laplacian of Gaussian values characterize clumps of bacteria

The most defining feature of bacterial clumps was the low, highly non-homogeneous signal in the total DNA channel. However, the standard deviation of brightness failed to distinguish them from other classes of ROIs. Another characteristic feature was the lack of well-defined borders, which was effectively captured by small values of the average Laplacian of Gaussian (LoG) of the total DNA channel.

#### Five features distinguish eight classes of ROIs

The progression of NET formation, the observed morphologies of different stages in fluorescent microscopy images, and the mathematical features that characterize them motivate the classification of ROIs into the following classes: *PMN neutrophils*, unstimulated, polymorphonuclear neutrophils, with a moderate cell size and brightness, and a lack of signal on the extracellular DNA channel; *RND neutrophils*, cells with decondensed chromatin and rounded nucleus, with higher brightness than PMN neutrophils; *NER neutrophils*, neutrophils with ruptured nuclear envelope, with larger cell sizes than RND neutrophils and possibly lower brightness than PMN neutrophils; *PMP neutrophils*, neutrophils with a permeabilized plasma membrane, with detectable signal on the extracellular DNA channel; *NETs*, Neutrophil Extracellular Traps, with an intense signal on the extracellular DNA channel, low to no signal on the total DNA channel, and noticeable standard deviation of the brightness; *Groups of bacteria*, with low brightness and LoG values in the total DNA channel, and possibly moderate signal on the extracellular DNA channel due to possible co-localization with NET fragments; *Autofluorescence artifacts* in the extracellular DNA channel, with a moderate, mostly homogeneous signal in this channel; *Unclassified ROIs* in the total DNA channel, corresponding to ROIs that do not match any of the previous classes, including neutrophil cells during a transition between the described classes. Note that a combinatorial approach to class characterization with ROI features allowed us to use a set of features smaller than the set of classes.

### 4.3 Classification workflow

The processing of a single fluorescent image with two channels (an extracellular DNA channel and a total DNA channel) is represented schematically in Fig. 5. In the pre-processing stage, the user may decide to use one of a number of filters provided by PartSeg (including the Gaussian and the median filter) on any or both channels. In the subsequent ROI detection stage, segmentation is performed on both channels by simple thresholding and detecting connected components. The brightness thresholds for both channels can be adjusted by the user during an interactive session of PartSeg to obtain a segmentation that matches a manual annotation.

**Figure 5:**
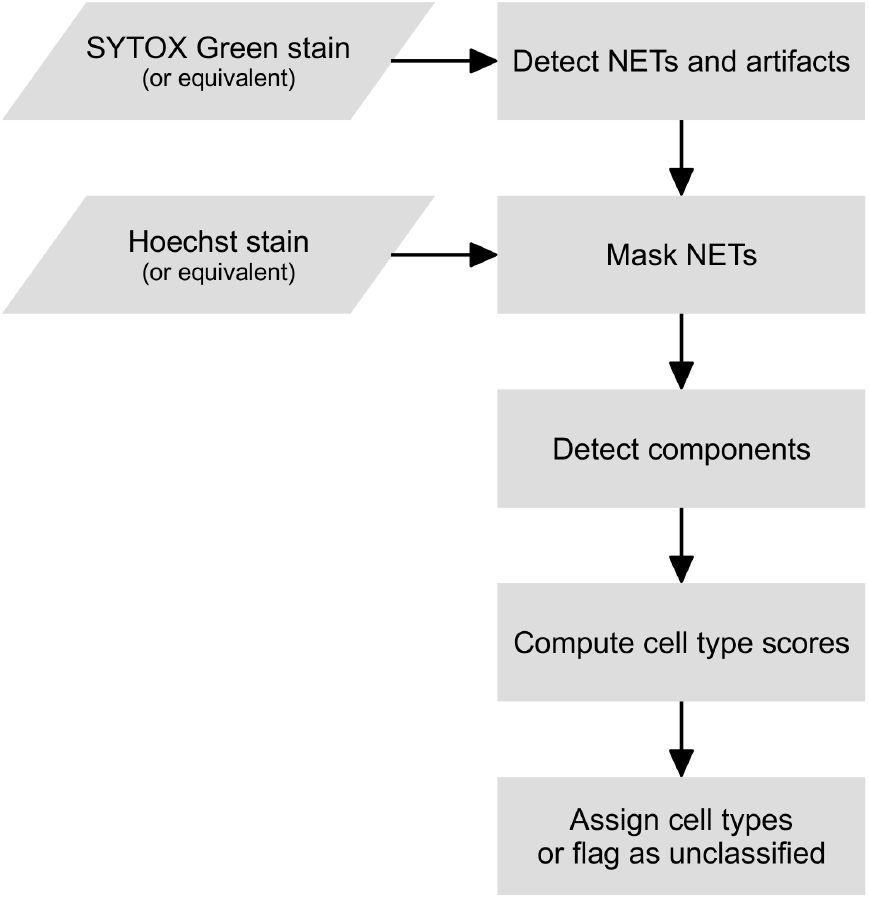
A flowchart of Trapalyzer.

In the first classification stage, Trapalyzer classifies ROIs on the extracellular DNA channel. Small ROIs on the extracellular channel, which typically correspond to neutrophils with a permeabilized plasma membrane, are not processed at this stage. If an ROI has a sufficient size and its average brightness and standard deviation are within user-defined ranges, it is classified as a NET. Otherwise, it is flagged as an “extracellular unknown” class, corresponding e.g. to autofluorescence artifacts.

In the second classification stage, Trapalyzer classifies ROIs in the total DNA channel. For each class, Trapalyzer computes a score that measures whether a given ROI matches its user-defined characterization. The general idea behind the class score is to ensure that all of the ROI features are within appropriate ranges for this class, with an error margin that allows some flexibility when defining the ranges.

Formally, let *x* be the value of a particular feature (e.g. brightness) of a given ROI, and let [*l, u*] be the acceptable interval for this feature for a given class of ROIs (e.g. NER neutrophils). For a single feature, we define a partial score function *S*(*x*; *l, u, s*), where the *s* parameter controls the extent of the error margins. The idea behind the partial score function is that *S*(*x*; *l, u, s*) equals 1 if *x* ∈ [*l, u*], and falls smoothly to 0 as *x* becomes distant from the interval [*l, u*], with the decrease rate controlled by *s*. Formally, we want *S* to be equal to 0 either when 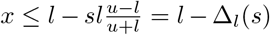 or 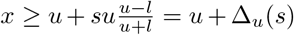.

This way, the left and right error margins for the characteristic range, Δ_*l*_(*s*) and Δ_*u*_(*s*), adjust to the interval length and its boundary values. Because of this, a single value of the error margin parameter *s* can be set for all features regardless of their units and typical values.

The properties described above are satisfied by the following function, where *l*_0_ = *l* − Δ_*l*_ and *u*_0_ = *u* + Δ_*u*_ define the range in which we want *S* to have a non-zero value:

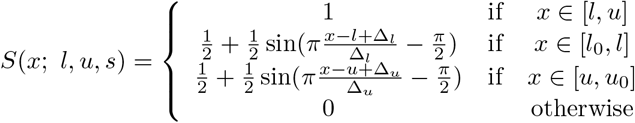

To obtain a final score for a given class, we multiply the partial scores for all the features of the analyzed ROI. This ensures that a ROI fully matches a class when all the features are within or sufficiently close to their acceptable ranges.

A ROI is determined as belonging to a given class if the score for this class is sufficiently high (by default above 0.8), and the scores for all the other classes are sufficiently low (by default below 0.4). The upper threshold for the scores of “competing” classes ensures that a ROI is classified unambiguously. If a ROI does not reach a sufficiently high score for any class, or reaches a high score for more than one class, it is flagged as an unknown class that requires a manual inspection.

During the second classification stage, we mask the regions occupied by NETs and extracellular artifacts and ignore ROIs in those regions. This is because, if a neutrophil lies within such a region, it is either difficult or impossible to accurately distinguish whether or not it has a permeabilized plasma membrane, and classification would therefore be unreliable. On the other hand, the number of such ROIs is typically small compared to the overall number of ROIs, and ignoring them had a relatively small influence on the overall performance of the software.

After both classification stages are completed, Trapalyzer evaluates the quality of image annotation. We define the quality score as *Q* = 100 · (1 − *U/S*)%, where *U* denotes the area covered by unclassified ROIs and *S* denotes the area covered by all detected ROIs. We use the areas of ROIs instead of their numbers to make the score robust to small artifacts that otherwise do not interfere with the analysis.

### 4.4 Validation on a benchmark data set

In order to assess Trapalyzer’s accuracy, we have compared it to previously reported results achieved with convolutional neural networks (CNNs) on a publicly available benchmark data set [14]. To match the original study, we have restricted the classification to four classes: PMN, NER and PMP neutrophils and NETs (the NER neutrophils were referred to as *decondensed* in the original work).

After tuning the parameters on a training set of 10 images, we have used Trapalyzer’s batch processing mode to analyze a validation data set consisting of 57 images. The resulting annotation was then compared with the manual one provided with the data set. We used an Intersection over Union (IoU) threshold of 0.10, meaning that we match objects if the overlap of their bounding boxes is at least 10% of their joint area. The IoU value was selected based on previous results in NET quantification [14, 9]. In case of more than two ROIs with overlapping bounding boxes, the pair with the highest IoU was selected as a match.

#### Consistent image acquisition conditions are crucial for automated image analysis

The results of Trapalyzer annotation are shown in Table 1. On average, 11.90% of ROIs were unclassified in each image, with two images exceeding 50% due to atypically low brightness of cells, likely caused by a low exposure time.

**Table 1:**
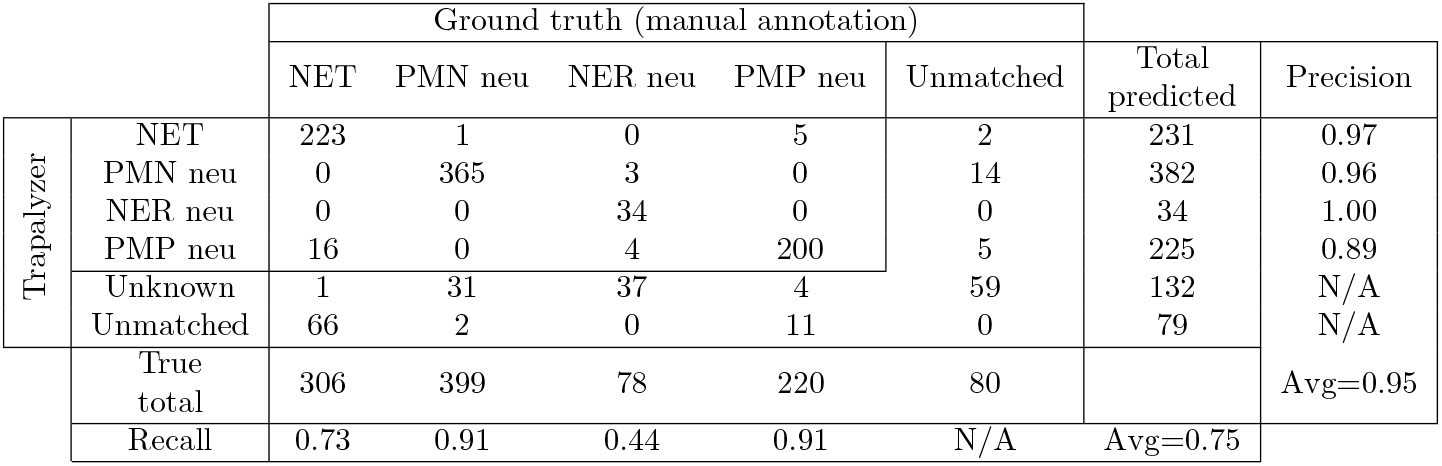
The results of an analysis of a publicly available benchmark data set of 57 fluorescent microscopic images.

|  |  | Ground truth (manual annotation) |  |  |  |  | Total predicted | Precision |
| --- | --- | --- | --- | --- | --- | --- | --- | --- |
|  |  | NET | PMN neu | NER neu | PMP neu | Unmatched |  |  |
| Trapalyzer | NET | 223 | 1 | 0 | 5 | 2 | 231 | 0.97 |
|  | PMN neu | 0 | 365 | 3 | 0 | 14 | 382 | 0.96 |
|  | NER neu | 0 | 0 | 34 | 0 | 0 | 34 | 1.00 |
|  | PMP neu | 16 | 0 | 4 | 200 | 5 | 225 | 0.89 |
|  | Unknown | 1 | 31 | 37 | 4 | 59 | 132 | N/A |
|  | Unmatched | 66 | 2 | 0 | 11 | 0 | 79 | N/A |
| True total |  | 306 | 399 | 78 | 220 | 80 |  | Avg=0.95 |
| Recall |  | 0.73 | 0.91 | 0.44 | 0.91 | N/A | Avg=0.75 |  |

#### Simple classification workflow achieves precision on par with convolutional neural networks

Over all ROI classes, Trapalyzer achieved an average precision of 95%, higher than 91% reported for the CNN classifier trained on 188 images. Precision varied slightly between classes, with the highest equal 100% for NER neutrophils, and the lowest equal 89% for PMP neutrophils, caused by annotating some NETs as neutrophils with permeabilized plasma membrane. Separating NETs from PMP neutrophils is difficult due to the gradual nature of chromatin release into the extracellular environment, so mismatches between the manual and automatic annotation are to be expected.

#### Imperfect parameter values decrease recall

Trapalyzer achieved an average recall of 75%, lower than 93% reported for the CNN. However, the recall varied greatly between classes, from 44% for NER neutrophils up to 91% for PMN and PMP neutrophils. Approximately half of the NER neutrophils were flagged as an unknown ROI class, suggesting imperfect parameter estimation from the training data set. As shown in the next section, in practical applications of Trapalyzer, this problem can be easily handled by re-tuning the parameters.

#### Quantifying NET area is more reliable than NET count

The recall value for neutrophil extracellular traps was 73%, caused by difficulties with matching Trapalyzer and manual annotations. Inspecting selected images showed that Trapalyzer missed fragments of NETs with a low brightness, causing low IoU values due to large differences between detected and manually generated bounding boxes. Moreover, NETs tend to merge if released by closely located cells and, although a human expert can detect such cases and identify individual nets, our approach to segmentation treats them as a single object. This agrees with observations made by other authors that the numbers and areas of individual NETs are difficult to quantify algorithmically, and quantifying the total image area covered by NETs is more reliable [9].

### 4.5 A case study of a neutrophil-*E. coli* co-culture

As an example application of Trapalyzer, we have performed a detailed analysis of the 120 fluorescent microscopy images of neutrophils incubated with or without *E. coli* bacteria, which we used to establish quantifiable classes of ROIs in the previous subsections. We have tuned the software’s parameters on a set of selected 10 images and further adjusted them on images with large numbers of unclassified ROIs. The correctness of annotation was then validated by an expert on 5 images. An example annotation of a fragment of an image is shown in Fig. 6.

**Figure 6:**
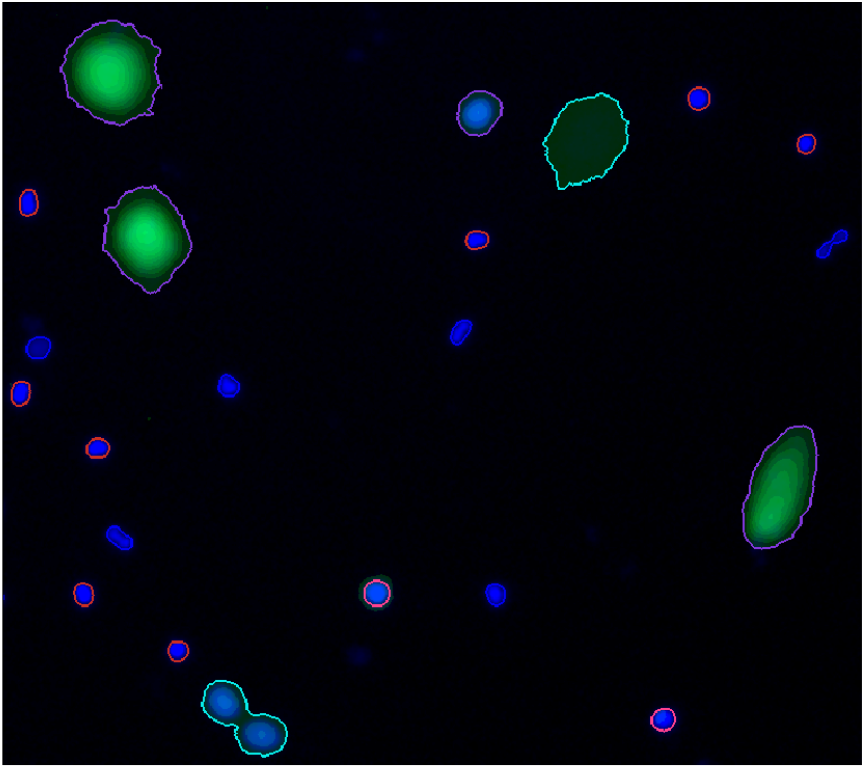
An example Trapalyzer annotation of a fragment of a fluorescent microscopy image. Our software identified four NETs (magenta outline), an autofluorescence artifact (turquoise outline), multiple PMN neutrophils (blue), RND neutrophils (red) and two PMN neutrophils (pink). Two emerging NETs in the lower left corner were misidentfied and flagged as a potential artifact for manual inspection.

#### Population-level results support the current model of NET formation

The total number of cells detected by Trapalyzer stayed approximately constant over the duration of the experiment (Supplementary Fig. S2), in agreement with the fact that only a few cells release NETs, showing that the experimental conditions and image acquisition methods were consistent. We observed a gradual decrease of the number of PMN neutrophils over time and the transition to RND, NER, and PMP cell morphology types (Supplementary Fig. S1), in agreement with the observations made for individual cells in [18]. NETs were formed continuously throughout the experiment and their number seemed to grow linearly in all experimental conditions, including the control group without bacteria (Supplementary Fig. S2). However, the rate of NET formation is higher in the co-cultures than in the control, indicating that the presence of bacteria successfully induced NET formation.

#### ROI-level results suggest an additional stage of NET formation

The properties of ROIs classified as neutrophils at different stages of NET formation are shown in Fig. 7. Different classes were clearly separated by the ROI features used in our classification workflow, which confirms their distinct natures. We observed a gradual increase in cell size as NET formation progressed. The average brightness was the highest for RND neutrophils, likely due to chromatin decondensation and increased dye affinity, and decreased after the rupture of the nuclear envelope when the chromatin occupied a larger area. A visibly bi-modal distribution of the brightness of RND neutrophils suggests that there may be an additional stage NET formation, which causes this group to be composed of two different types of cell morphologies. This phenomenon requires further studies.

**Figure 7:**
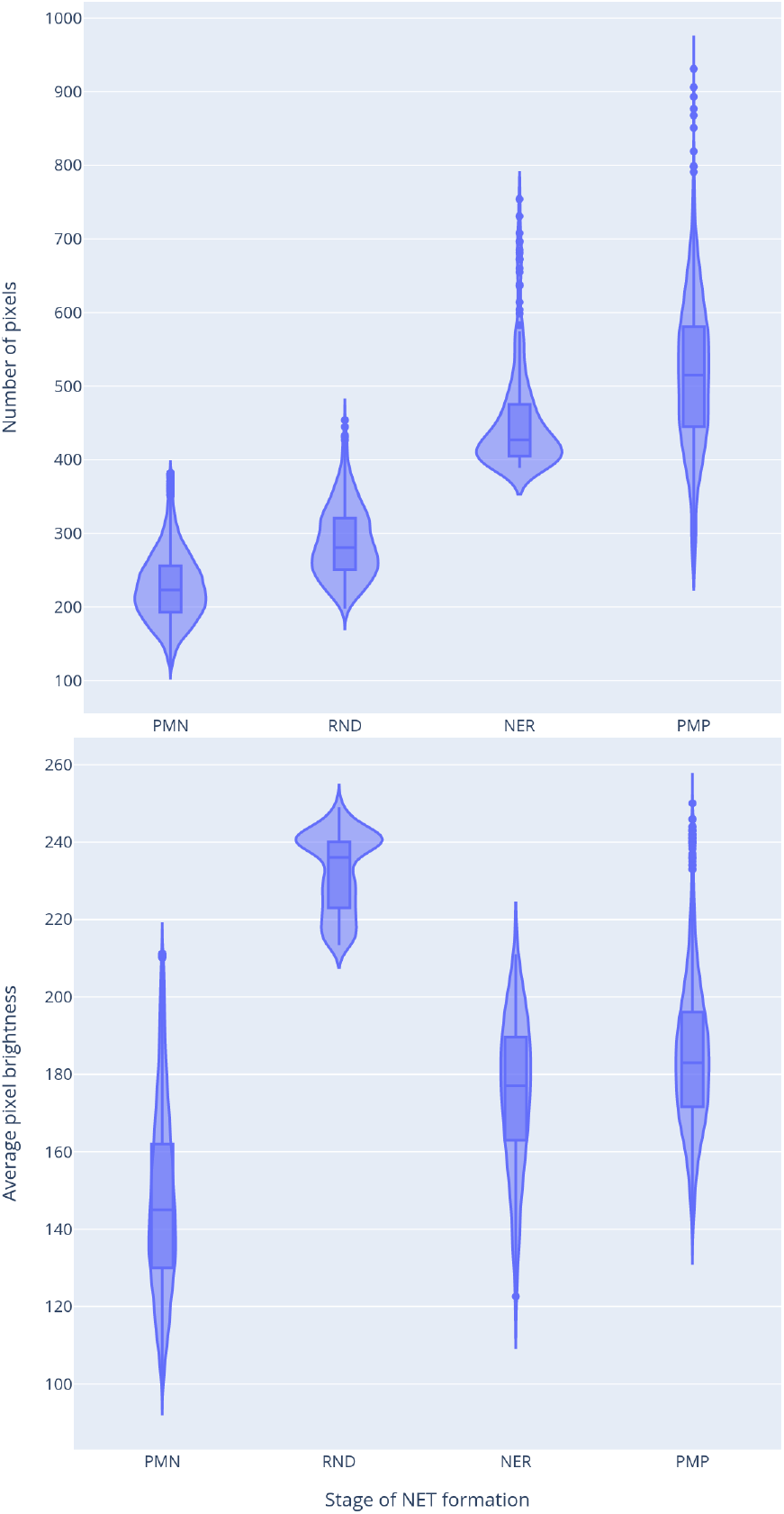
Properties of ROIs classified as neutrophils at different stages of NET formation. Top: ROI area (in number of pixels). Bottom: average pixel brightness in the total DNA channel. The third feature, average brightness on the extracellular channel, had a non-zero value only for the PMP neutrophils.

#### Mathematical modelling of the dynamics of a neutrophil population is challenging

Neutrophils stayed viable for a prolonged period of time in the negative control without bacteria. From the perspective of mathematical modeling, this means that traditional models based on ordinary differential equations may not be suitable to describe an *in vitro* cultured population of neutrophils. More sophisticated mathematical techniques, such as delay differential equations, may be required to model the dynamics of NET formation in such experiments.

#### A small number of false positives did not influence the overall conclusions

The number of bacterial groups grew exponentially in time when bacteria were present in the sample (Supplementary Fig. S2). This suggests that, in our experimental conditions, neutrophils had a limited capability to eliminate pathogens. We have observed a small number of false positive results in the control group without bacteria. In 8 out of 40 images for this experimental condition there were between one and three improperly detected clumps of bacteria, corresponding to image artifacts and misclassified neutrophil cells. In comparison, in the experimental condition with four bacterial cells per neutrophil, there were up to 100 bacterial groups for t=180 min. As a consequence, the number of false positive detections of bacterial clumps was comparatively small and did not influence the overall conclusions.

## 5 Discussion and conclusions

Software tools designed for the analysis of microscopic images of neutrophils and neutrophil extrcellular traps (NETs) can be roughly partitioned into two groups. The first group consists of tools based on machine learning, such as convolutional neural networks (CNNs) or support vector machines (SVMs). The second group consists of tools based on classical image processing techniques, such as edge detection, image filtering etc.

Trapalyzer employs the latter approach. It detects ROIs using a thresholding operation and classifies them as NETs, neutrophils at different stages of NET formation, or clumps of bacteria based on a handful of features such as area and brightness. An ROI is assigned to a class if it matches it unambiguously according to a scoring function.

Our reasons to make Trapalyzer machine learning-free are twofold. First, machine learning algorithms require extensive training data sets on which they learn how to distinguish between different types of objects. Preparing such a data set can take many days for a trained expert. The algorithm is then capable of classifying detected objects in a test data set only as long as their morphology resembles the objects encountered in the training one. This is particularly limiting in case of NETs, because their morphologies may differ depending on the experimental conditions, such as the substance used to stimulate neutrophils to release the traps [3]. Even minor changes in experimental conditions, such as microscope magnification or exposure time, may require to re-train the algorithm. This negates one of the purposes of such tools, which is to make analyses easier, faster, and less laborsome. Still, NETs retain certain characteristic features, such as a large size and fluorescence in the presence of cell-impermeable DNA dyes, that can be used to detect and quantify them with a high degree of accuracy.

The second reason to avoid the use of machine learning algorithms is that NET formation is not yet a fully understood phenomenon, especially on the cellular level [18]. A lot of experimental work is still required to characterize different stages of NET formation and cell morphologies at those stages. Because of this, our goal with Trapalyzer was to offer a tool which helps the biologists, instead of replacing them. In this context, the advantage of image processing techniques is that they allow for a far greater degree of control over the classification process and interpretability of results. The user may freely decide which features to use for classification, which cell types are of interest in a given experiment, and specify the appropriate ranges of parameters.

Trapalyzer signals which ROIs are difficult to classify, allowing the user either to classify them manually, to adjust the program’s parameters, or to simply ignore them if appropriate. Reporting such ROIs by the software is of a particular importance when the DNA is stained with a green dye, such as SYTOX™ Green, as the autofluorescence on the green channel may cause artifacts in the images [2]. Detecting artifacts allows Trapalyzer to detect images for which the annotation may be unreliable, and highlight regions of the images which should be analyzed manually. This way, artifacts and atypical cell morphologies can be easily detected and handled.

One of the main contributions of this work is the identification of a set of just four features that allow us to classify ROIs to a high degree of accuracy. Because of this, Trapalyzer requires just a handful of user-specified parameters to perform the classification. Once set, the parameters apply as long as image acquisition parameters remain constant. Since the parameters depend on the equipment used in a particular laboratory, we do not include a default set of parameters in Trapalyzer, but provide an easy to follow tutorial with a step-by-step procedure of tuning them, available on the project website.

In this work, we detect and classify polymorphonuclear neutrophils solely based on their size and average brightness. Due to their characteristic shapes, some measure of ROI circularity could potentially also be a characteristic feature of this morphology. Two common measures of this feature are the ratio of the ROI area to squared perimeter and the ratio of the ROI area to squared diameter. However, none of those measures was capable of distinguishing polymorphonuclear neutrophils from other stages of NET formation and improve the accuracy of classification. This is because some segments corresponding to those cells are elongated but otherwise highly regular, and both circularity measures are high in such cases. We did not find any mathematical characterization of the irregular shape of polymorphonuclear neutrophils that would be useful for our purposes.

To our knowledge, Trapalyzer is the only currently available computer program capable of quantifying not only NETs in terms of their number and area, but also the numbers of neutrophils at different stages of NET formation in experiments where extensive training data sets are not available.

## 5.3 Author’s Contributions

MAC and AG conceived the study. MAC, GB and AMH designed the software. GB implemented the software. MAC and AMH designed the co-culture experiment. AMH and WK performed the experiment. MAC analyzed the experimental data and wrote the draft manuscript. AG and UD supervised the work. All authors have read and approved the manuscript.

## 6 Supplementary figures

**Figure S1:**
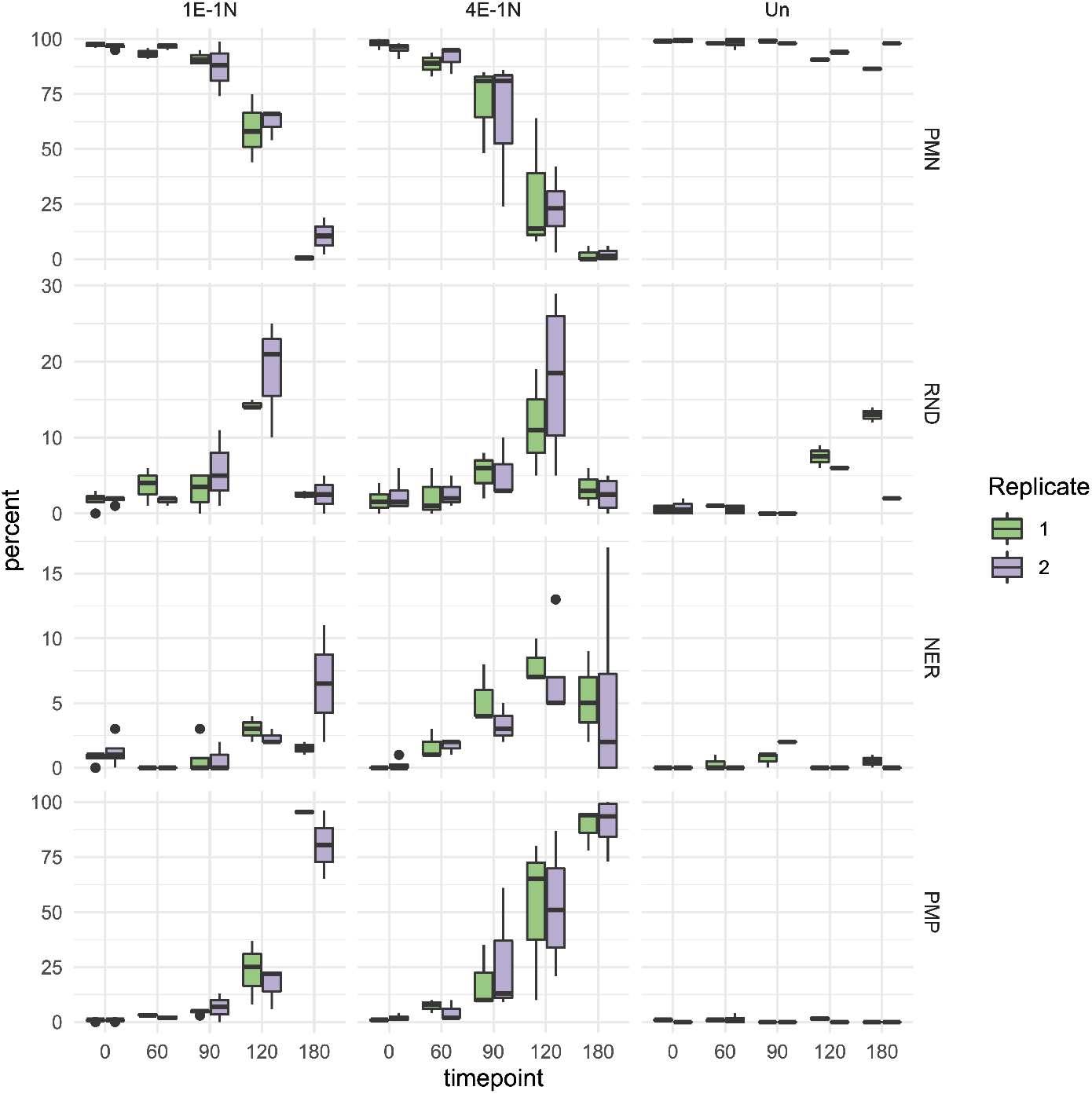
The progression of NET formation in a neutrophil-*E. coli* bacteria co-culture experiment. Each row shows the percentage of cells in a given stage of NET formation. Data was obtained by analyzing 120 fluorescent microscopy images with Trapalyzer. Four images were taken for each timepoint, experimental condition, and technical replicate. Images with low annotation quality were discarded from the results.

**Figure S2:**
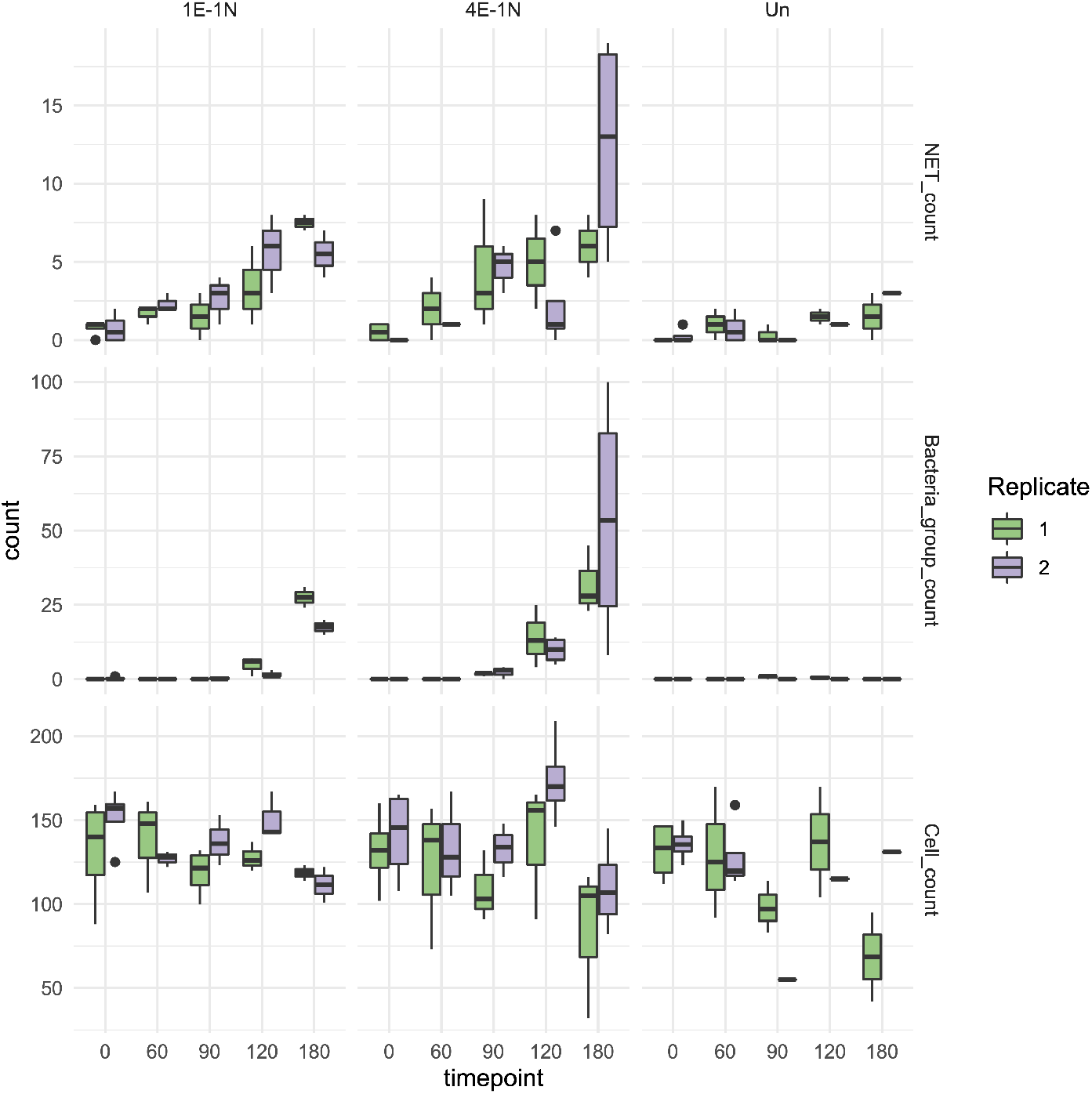
The numbers of neutrophil extracellular traps, groups of bacteria, and the total number of neutrophil cells at all stages of NET formation, obtained by analyzing fluorescent microscopy images from the neutrophil-*E. coli* co-culture experiment with Trapalyzer. Images with low annotation quality were discarded from the results.

